# Combining Multiscale Diffusion Kernels for Learning the Structural and Functional Brain Connectivity

**DOI:** 10.1101/078766

**Authors:** Sriniwas G. Surampudi, Shruti Naik, Avinash Sharma, Raju Surampudi Bapi, Dipanjan Roy

## Abstract

The activation of the brain at rest is thought to be at the core of cognitive functions. There have been many attempts at characterizing the functional connectivity at rest from the structure. Recent attempts with diffusion kernel models point to the possibility of a single diffusion kernel that can give a good estimate of the functional connectivity. But our empirical investigations revealed that the hypothesis of a single scale best-fitting kernel across subjects is not tenable. Further, our experiments demonstrate that structure-function relationship across subjects seems to obey a multi-scale diffusion phenomenon. Based on this insight, we propose a multiple diffusion kernel model (**MKL**) along with a learning framework for estimating the optimal model parameters. We tested our hypothesis on 124 subjects’ data from publicly available NKI_Rockland database. The results establish the viability of the proposed model and also demonstrate several promising features as compared to the single kernel approach. One of the key strengths of the proposed approach is that it does not require hand-tuning of model parameters but actually learns them as part of the optimization process. The learned parameters may be suitable candidates for future investigation of their role in distinguishing health and disease.

## 1 Introduction

’Connectome’, coined after ’Genome’ that stands for the effort of mapping genetic code, refers to the map of the brain’s neural connections ([1], [2]). Consequently ’Connectomics,’ like Genomics, concerns with the assembling and analyzing connectome data sets. Two primary classes of connections are often considered in this analysis – the anatomical links via fiber pathways (referred to as the structural connectivity **SC**) and the statistical dependency measures such as correlation of activity in distinct anatomically segregated regions (designated as the functional conncectivity **FC**). After the accidental discovery of slow correlated fluctuations in the resting state functional magnetic resonance images (**rs-fMRI**) of the brain by Biswal’s group [3], investigation of these task-independent, spontaneously evoked cortical responses known as the resting state brain activity gathered momentum. Functional connectivity analysis of the resting-state brain activity led to the discovery of groups of brain regions exhibiting correlated activation such as the default mode network (DMN), vision, auditory, language, sensori-motor, salience, executive, and attentional networks ([4],[5]).

The organization of brain networks has been gainfully captured using graph representation, with nodes representing the brain regions and the edges denoting the structural or functional connections between them. A large number of graph theoretic measures such as centrality, modularity, small world properties, etc have been borrowed for characterizing the topological properties of brain networks. A brain connectome with *n* cortical regions can be captured as an *n × n* matrix, *C*. Here the matrix entries *C*_*ij*_ encode the connection strength between two regions *i* and *j*. In **SC** matrix, the entries often stand for density of white matter fiber tracts extracted from a diffusion weighted image such as the diffusion tensor image (**DTI**). On the other hand in the **FC** matrix, these entries often denote the Pearson correlation coefficient computed between the blood oxygen level dependent (BOLD) changes in the resting state activity of the two brain areas. Another aspect of brain graphs that is important is its size. The size of the network, i.e., the number of nodes or regions of interest (**ROI**s), is determined by how the brain is parcellated into regions. High resolution parcellations provide more information about both anotomical structures as well as their functional behavior.

The relation between structural and functional connectivity in health and disease is being investigated intensely (see for example, [7], [8], [9], [10]). If there is a direct anatomical connection between brain regions, it is often observed that these regions are also functionally connected. However the converse is not necessarily true. The question of how SC shapes FC has been the object of computational modeling but remains an open question. Whole-brain computational models with the connectivity constrained by SC matrix have been employed to generatively model the FC matrix (e.g., [11], [12]). Recently, a linear graph-theoretic dynamic model has been proposed that captures the structure-function relationship using a simple, low-dimensional network diffusion model [6]. While the proposed diffusion model with a single scale was sufficient to capture the long-range correlation structure of brain activity and the results seem to indicate better correspondence than was possible with other non-linear models (see for example, [13]), there are critical shortcomings of this model.

Their results suggested the possibility of a single-scale of diffusion that enables maximum correlation between observed **FC** and estimated **FC** in experiments with eight subjects. Consequently we investigated the viability of this hypothesis over a larger subject pool. In our simulation experiments we found that the diffusion scale for maximal correlation (between the empirical and observed **FC**s) varies widely across subjects (see Figure 1). On the other hand, we also observed that for an individual subject, diffusion at different scales reveals multiscale relationships among various ROIs. Thus, multiple scale dependent diffusion kernels over **SC** can be interpreted as components of **FC** at different scales (see Figure 2). Interestingly, similar multiscale behavior of diffusion kernel has been exploited in other research domains such as Computer Vision [14, 15]) and Robotics [16]. This naturally motivated us to formulate a multiple diffusion kernels derived from the **SC** and a learning scheme that optimizes the linear combination of the multiple kernels that best fit the observed **FC**s.

**Figure 1:**
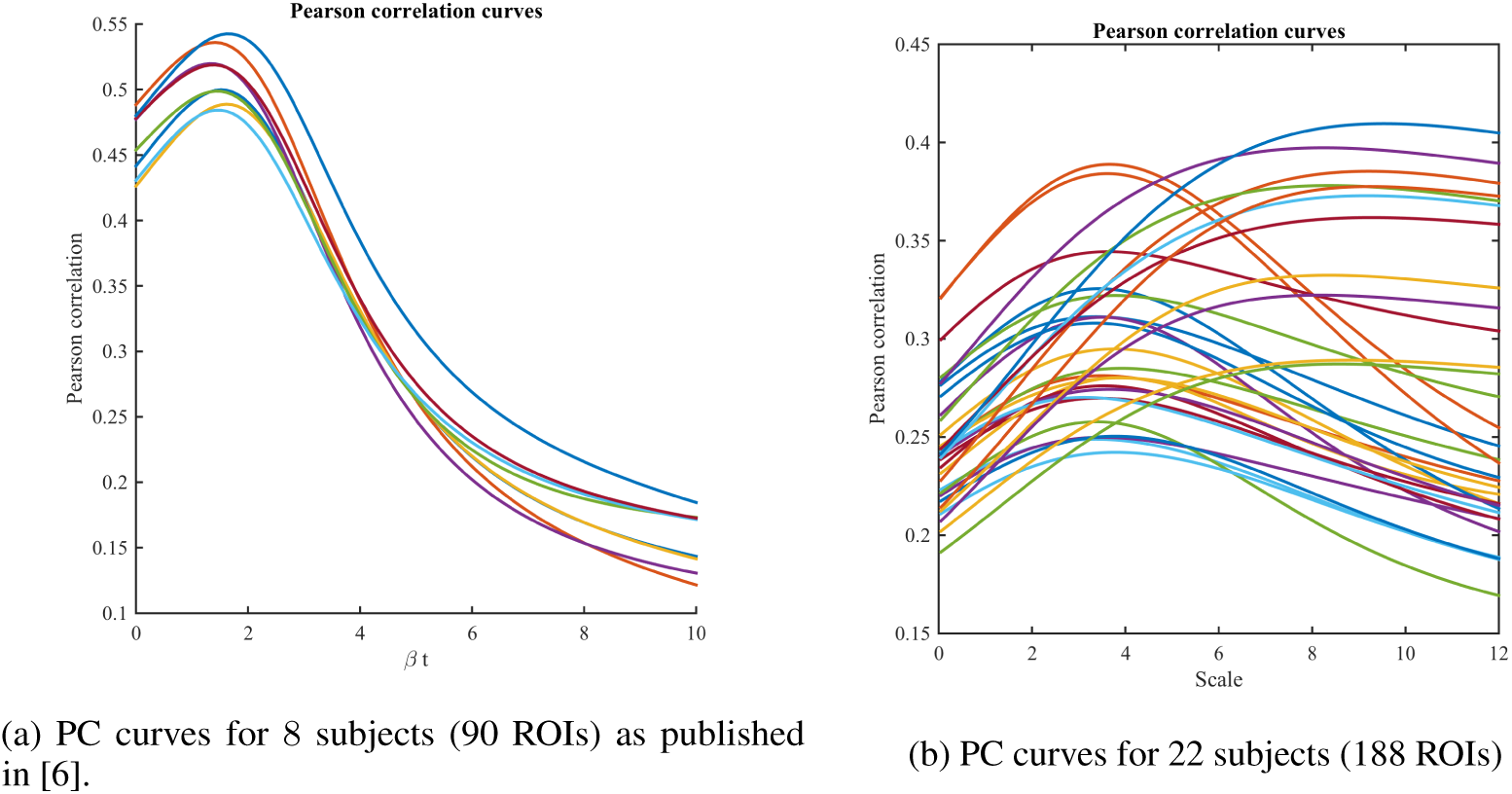
(a) Pearson Correlation (PC) curves for smaller dataset with smaller parcellation exhibit a single optimal scale of diffusion. (b) However, for a bigger dataset with larger parcellation, the results suggest that no single optimal scale exists.

**Figure 2.**
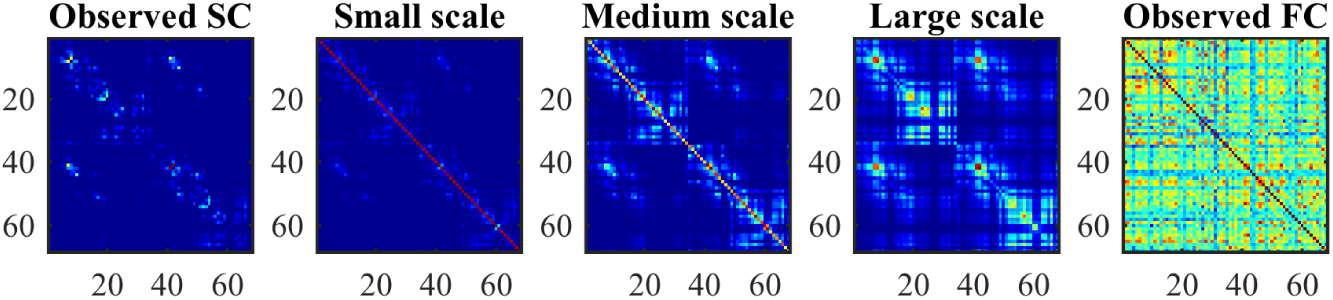
Diffusion at multiple scales for a single subject. The results of diffusion over **SC** at three different scales are shown in the three middle panels. It appears that a linear combination of these multiscale diffusion matrices would approximate the observed **SC**-**FC** relationship better.

In this paper, we propose a Multiple Kernel Learning (**MKL**) formulation for reconstructing **FC**s by finding optimal parameters that linearly combine diffusion kernels at different scales. MKL techniques are well explored in ML community [17, 18]. Our proposed MKL model while retaining the parsimony of a simple linear approach, proposes a novel learning scheme for optimizing the best-fitting kernels for **SC-FC** mapping. It is important to note that here we learn the coefficients of linear combination of multiple diffusion kernels (defined at different scale). Therefore, the resultant kernel need not be interpreted as a diffusion kernel at some optimal scale. Our detailed empirical results demonstrate the validity of the proposed MKL model on a larger dataset. In the next section, MKL formulation is outlined, followed by the presentation of the results.

## 2 Proposed Method

We extend the linear graph-theoretic dynamic model proposed in [6] for learning the structure-function relationship using a novel MKL formulation.

Let **W** be the weighted adjacency matrix representing the structural connectivity (**SC**). The symmetric Laplacian matrix can be obtained from **W** as: 
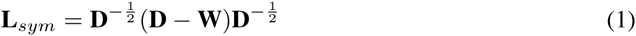
where, **D** is the degree matrix of **W**. The scale dependent diffusion kernel matrix is computed as:

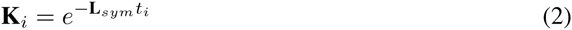

As diffusion kernels transform **SC** into nonlinear spaces, we hypothesize that the linear combination of these nonlinear mappings would give rise to a good estimation of **FC** (see Eq.3).

Let *τ* = {*t*_1_…, *t*_*m*_} be the set of *m* diffusion scales. For each *t*_*i*_ a corresponding **K**_*i*_ is obtained using Eq. 2. Let *α* = {*α*_1_…, *α*_*m*_} be the set of coefficients of linear combination (also called here as mixing coefficients) for the corresponding *m* kernel matrices. These mixing coefficients are subsequently learned while solving an optimization formulation that minimizes the squared error between empirical **FC** (provided as ground truth) and predicted **FC** (see Eq. 3).

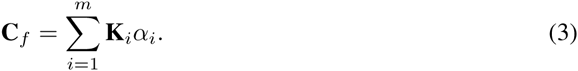

Let the training set size be *p*, i.e., the number of subjects for whom **SC-FC** pair is considered during the training phase. We can write the optimization function as:

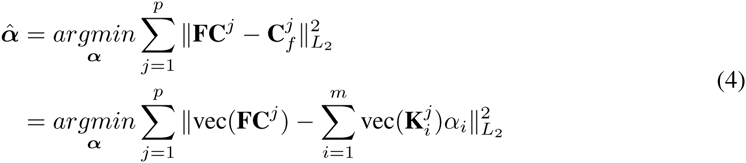

Here, vec(*⋅*) converts an *n × n* matrix into an *n*^2^ *×* 1 vector.

Let 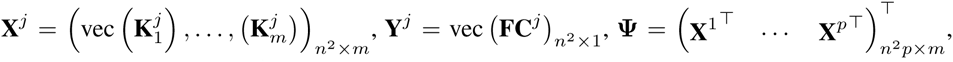 and 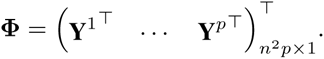. Then,

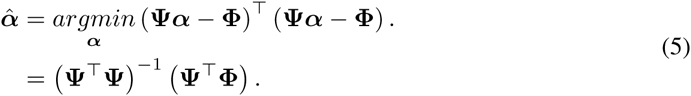

We find the least squares solution and divide ***α***ˆ by its sum to normalize the values.

In summary, the key differences between the single diffusion kernel formulation ([6] and the proposed MKL approach are the following:

- It can be noticed that we use multiple diffusion kernels (see equation 3) aiming to capture the multiscale relation observed in our empirical experiments as shown in Figures 1b and 2.
- Instead of empirically hand-picking the optimal mixing coefficients, we actually estimate them by minimizing an optimization formulation (see equation 5).

## 3 Results

### 3.1 Datasets

In order to investigate the behavior of the single diffusion kernel model of [6], we used **SC-FC** data of 22 subjects randomly chosen from the NKI_Rockland data [19]. The simulation results are shown in Figure 1b.

For further experiments on evaluating the performance of the proposed MKL model, we utilized data from a total of 124 subjects from the same NKI_Rockland database referred above. Of these 124, a randomly chosen half (62 subjects’ data) was used for training and the other half for testing. The results of these experiments are shown in Figures 3, 4, and 5.

### 3.2 Experiment & Results

In the first experiment, we compared the results from single kernel [6] versus our proposed MKL model. We use the Pearson correlation [11, 6] for comparing observed **FC** and predicted **FC**. In the single kernel case, we are directly picking the best kernel and computing the PC values for the test subjects. On the other hand, for the MKL model, parameters are estimated from training data and PC values are computed for each of the remaining 62 test subjects. Figure 3 shows the comparative results.

**Figure 3:**
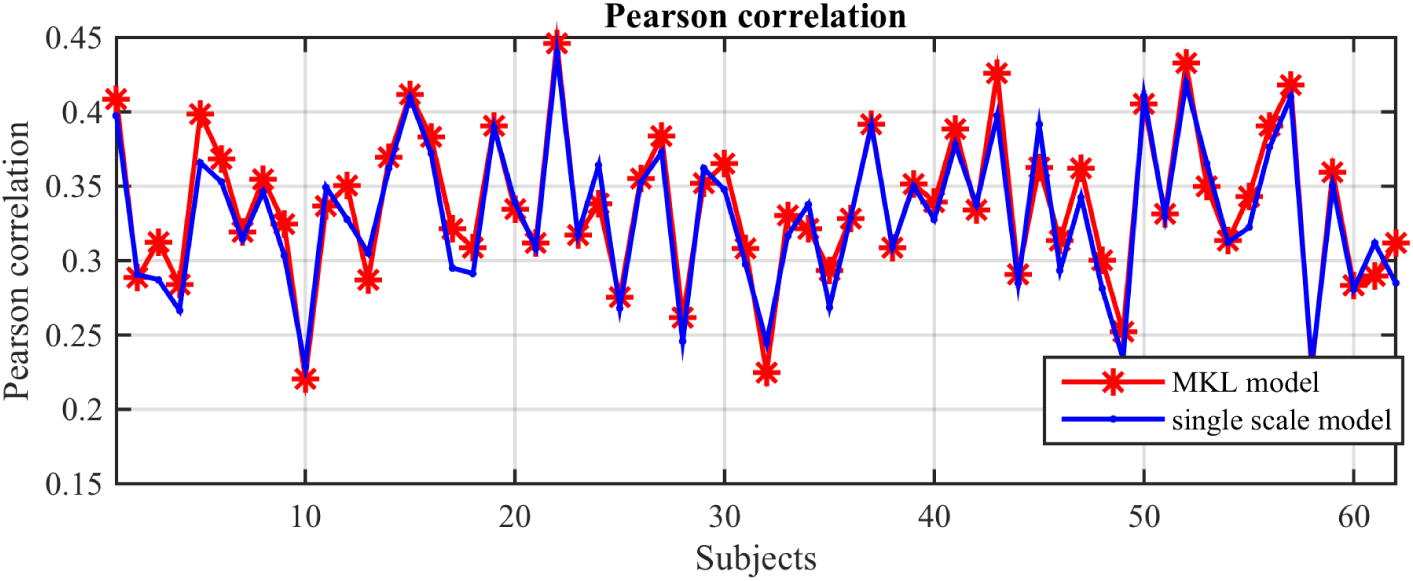
Comparison of the proposed MKL model versus the single scale model. As can be seen, the MKL model performs at par or better than the single scale model. It is to be noted that optimal scale for the single scale model was selected on the test subjects’ data by searching individually. On the other hand, the optimal parameters for the MKL model were learned through the optimization process on training data.

The next experiment demonstrates the robustness of the proposed method w.r.t. the choice of the number of scales i.e., parameter *m* (see also Equations 3, 4). Here we experimented with 5 different values of *m* (using the same train and test set used in the first experiment) and we can see in Figure 4 that the model performance is relatively stable (curves closely follow each other). As shown in Table 1 the performance of our method shown for five randomly chosen subjects improves with increasing values of *m* till *m* = 8 and then stabilizes. This was indeed the case with all other subjects. The increase in performance (from *m*=2 to *m* = 8) is attributed to the fact that with higher multiscale resolution our MKL model perhaps leads to a better reconstruction of the observed FC.

**Figure 4:**
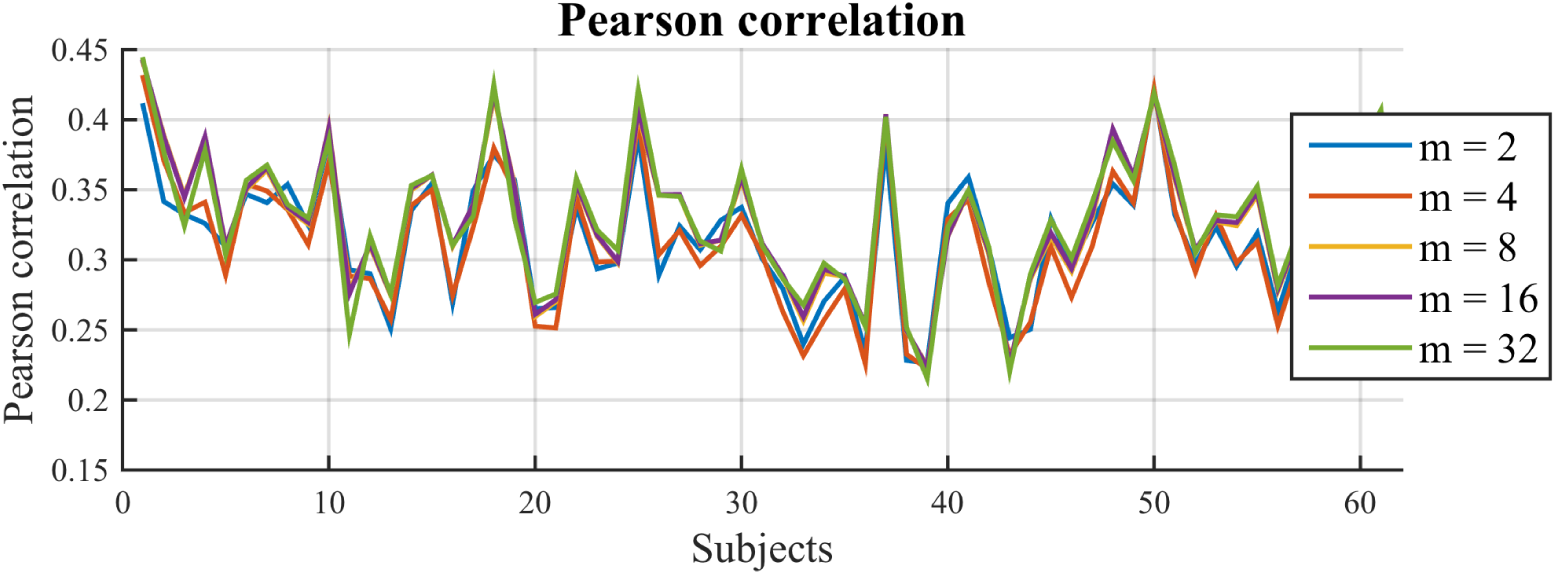
Performance stability of the proposed MKL model for different values of number of scales parameter.

**Table 1:** Pearson correlation w.r.t. different number of scales (parameter *m*).

| $m$ | Subjects | | | | |
| --- | --- | --- | --- | --- | --- |
|  | sub 1 | sub 2 | sub 3 | sub 4 | sub 5 |
| 02 | 0.34 | 0.24 | 0.28 | 0.28 | 0.29 |
| 04 | 0.41 | 0.32 | 0.28 | 0.34 | 0.30 |
| 08 | 0.44 | 0.34 | 0.31 | 0.35 | 0.32 |
| 16 | 0.43 | 0.43 | 0.31 | 0.35 | 0.32 |
| 32 | 0.43 | 0.43 | 0.31 | 0.35 | 0.32 |

In the final experiments, we try different configurations of scale values (i.e., ***τ***) for *m* = 10. Table 2 shows the values for 6 different scale configurations. Figure 5 shows the corresponding plot of PC curves. We can clearly see that these PC curves are highly overlapping thereby suggesting that the choice of scale values does not affect the performance of our proposed MKL model.

**Table 2:** Various set of scales ***τ***

| Scale | Configurations |  |  |  |  |  |
| --- | --- | --- | --- | --- | --- | --- |
|  | conf 1 | conf 2 | conf 3 | conf 4 | conf 5 | conf 6 |
| $t_1$ | 25.17 | 5.27 | 4.48 | 4.18 | 4.04 | 3.86 |
| $t_2$ | 11.64 | 2.43 | 2.10 | 1.94 | 1.87 | 1.79 |
| $t_3$ | 8.08 | 1.70 | 1.44 | 1.35 | 1.30 | 1.24 |
| $t_4$ | 5.95 | 1.25 | 1.06 | 0.99 | 0.95 | 0.92 |
| $t_5$ | 4.42 | 0.93 | 0.79 | 0.74 | 0.71 | 0.67 |
| $t_6$ | 3.22 | 0.67 | 0.57 | 0.54 | 0.51 | 0.50 |
| $t_7$ | 2.24 | 0.47 | 0.40 | 0.37 | 0.35 | 0.34 |
| $t_8$ | 1.41 | 0.30 | 0.25 | 0.24 | 0.23 | 0.21 |
| $t_9$ | 0.70 | 0.14 | 0.12 | 0.12 | 0.11 | 0.11 |
| $t_{10}$ | 0.05 | 0.12 | 0.01 | 0.01 | 0.01 | 0.01 |

**Figure 5:**
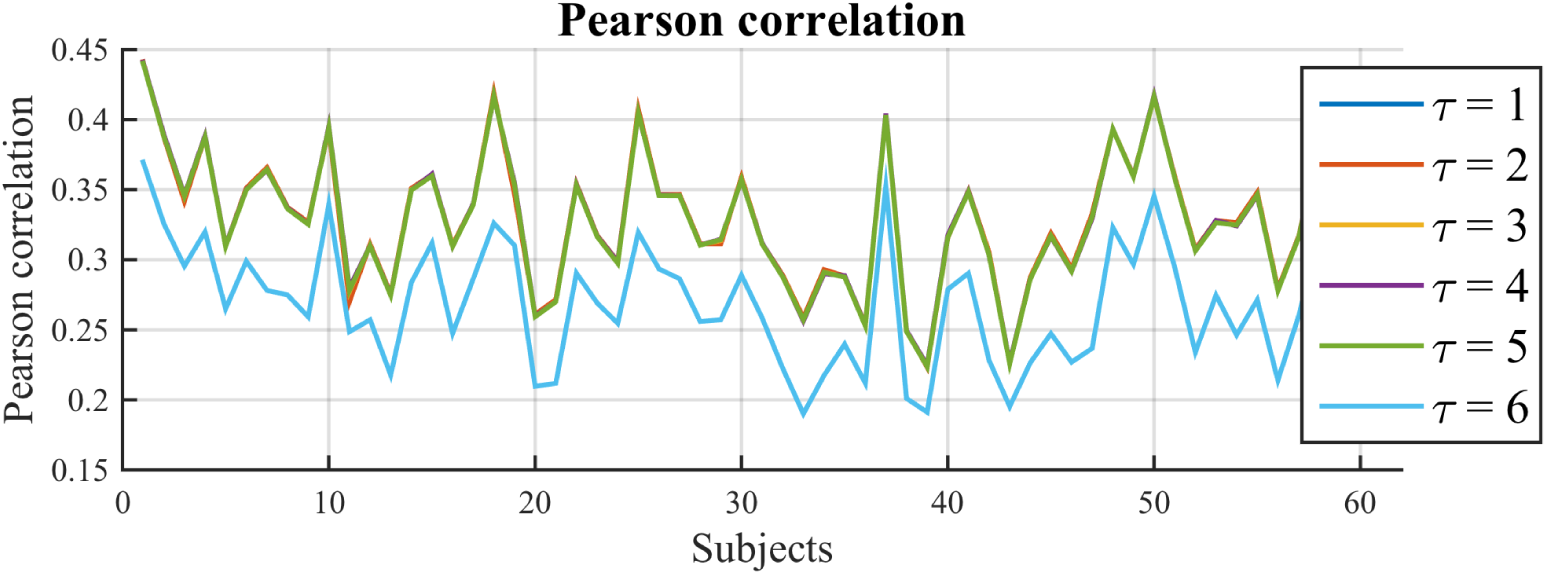
Performance of the MKL model across different scale sets or scale configurations. It is observed that the model performance is robust with respect to change of configurations.

## 4 Conclusion & Future Work

Inferring resting state functional connectivity **FC** from the underlying structure of the brain (**SC**) is a challenging open problem in computational neuroimaging. There are broadly two categories of models that attempt this problem – predictive versus generative. Simplistic network communication based models ([20], [21]) as well as diffusion kernel models fall in the former category. A range of generative models have also been proposed to this end using neuro-biologically detailed dynamical models ([22], [23]). Even though each model predicts **FC** fairly well, the onus of finding the right balance between biological realism and computational tractability remains on the modeler. Recently an elegant tractable dynamical model has been proposed by [6] which assumes linear dynamics between network nodes and predicts **FC** based on the assumption of macroscopic interactions of brain activity. However, our observation based on the simulation of the same model on finer parcellation of **SC** (more **ROI**s) suggests that the hypothesis of a single best-fitting diffusion kernel applicable across subjects seems to be not viable. This led us to formulate a more detailed **MKL** model, which obviates the need for manually searching for the single optimal kernel. More importantly, MKL model involves *learning* of the **SC-FC** mapping parameters on training set which can be directly used on unseen individual test subjects.

The results demonstrate that the proposed method predicts individual (subject-specific) **FC** at par or better than the previous model. Further, the model also has the power to explain inter-individual variability in **SC-FC** relationships. This possibly suggests the existence of latent variables across subjects, which when learnt, can explain the **SC-FC** relationship. This would in turn avoid the need for generating the long time course signals as is the case with more detailed computational models. Such machine learning approaches with the promise of simplicity might relieve the modelers from cumbersome subject-specific parameter tuning or an educated guess of parameters and initial conditions that are required in other non-linear dynamical models that are designed with a range of free parameters.

As part of future work, it will be interesting to find a better interpretation of the latent variables ***α***s and associated kernel matrices. Additionally, we would like to automatically discover the optimal scale configurations instead of assuming them to be chosen a priori. Further, more complex optimization formulations will be explored in order to improve the performance of the SC-FC mapping.

